# The role of *Cdx2* as a lineage specific transcriptional repressor for pluripotent network during trophectoderm and inner cell mass specification

**DOI:** 10.1101/124438

**Authors:** Daosheng Huang, Xiaoping Han, Ping Yuan, Amy Ralston, Lingang Sun, Mikael Huss, Tapan Mistri, Luca Pinello, Huck Hui Ng, Guocheng Yuan, Junfeng Ji, Janet Rossant, Paul Robson, Guoji Guo

## Abstract

The first cellular differentiation event in mouse development leads to the formation of the blastocyst consisting of the inner cell mass (ICM) and an outer functional epithelium called trophectoderm (TE). The lineage specific transcription factor CDX2 is required for proper TE specification, where it promotes expression of TE genes, and represses expression of *Pou5f1* (OCT4) by inhibiting OCT4 from promoting its own expression. However its downstream network in the developing early embryo is not fully characterized. Here, we performed high-throughput single embryo qPCR analysis in *Cdx2* null embryos to identify components of the CDX2-regulated network *in vivo*. To identify genes likely to be regulated by CDX2 directly, we performed CDX2 ChIP-Seq on trophoblast stem (TS) cells, derived from the TE. In addition, we examined the dynamics of gene expression changes using an inducible CDX2 embryonic stem (ES) cell system, so that we could predict which CDX2-bound genes are activated or repressed by CDX2 binding. By integrating these data with observations of chromatin modifications, we were able to identify novel regulatory elements that are likely to repress gene expression in a lineage-specific manner. Interestingly, we found CDX2 binding sites within regulatory elements of key pluripotent genes such as *Pou5f1* and *Nanog*, pointing to the existence of a novel mechanism by which CDX2 maintains repression of OCT4 in trophoblast. Our study proposes a general mechanism in regulating lineage segregation during mammalian development.

## INTRODUCTION

Totipotency, or the ability to form both embryonic and extra-embryonic tissues, is lost upon the first lineage segregation during mouse early embryonic development. As the blastocyst forms, cells that will become trophoblast are separated from the pluripotent cells of the embryo (Ralston and Rossant, 2005; Rossant and Tam, 2009). At this crossroad, cells decide whether to establish or repress pluripotency, in establishing the inner cell mass (ICM) and trophectoderm (TE), respectively. Embryonic stem (ES) cells originate from the inner cell mass (Evans and Kaufman, 1981; Martin, 1981), while trophoblast stem (TS) cells are derived from trophectoderm (Tanaka et al., 1998). These stem cell lines provide expandable yet pure cell populations for genome-wide analyses of gene regulatory mechanisms (Chen et al., 2008; Kidder and Palmer, 2010; Kim et al., 2008; Loh et al., 2006). Studies with these stem cell lines have illuminated our understanding the genetic networks that regulate the segregation of the first two lineages in the mouse pre-implantation embryo.

The transcription factor CDX2 acts early during the blastocyst formation, playing an instructive role in the formation of trophoblast. Loss of *Cdx2* in the embryos leads to ectopic expression of pluripotency markers in the TE (Strumpf et al., 2005), and over-expression of *Cdx2* in ES cells is sufficient to direct the formation of TS cells (Niwa et al., 2005). How CDX2 achieves its role via transcriptional regulation is therefore a central question. Nishiyama et al. characterized the genome-wide early responsive CDX2 targets when *Cdx2* was overexpressed in ES cells (Nishiyama et al., 2009a), and could not demonstrate direct binding of CDX2 to the regulatory regions of pluripotency genes. Rather, CDX2 interfered with a pro-pluripotency transcriptional complex during the early stages of CDX2 over-expression (Nishiyama et al., 2009a). However, the long-term activities of CDX2 in maintaining cell fate, in stem cell lines and *in vivo*, have not been extensively characterized.

Given the importance of understanding CDX2 targets in a biologically relevant setting, direct examination of CDX2 function in the embryonic TE tissues is needed. We first analyzed global gene expression in the TE of wild type embryos by mRNA-seq. We then developed micro-genomic methodologies to profile gene expression in individual *Cdx2* knockout blastocysts. We performed CDX2 ChIP-seq in TS cells, which identified CDX2 targets relevant to TE biology. Finally, we defined lineage-specific silencer regulatory regions that possess unique chromatin features, on a genome-wide level. Ultimately, we have integrated these data to present a holistic model of how CDX2 regulates the ICM/TE lineage segregation during mouse embryo development.

## RESULTS

### Comparison of *in vitro* trophoblast stem cell lines and **in vivo** trophectoderm progenitors

TS cells derived from blastocysts or Cdx2-overexpressing ES cells provide a useful platform to investigate gene regulatory networks of early cell commitment *in vitro* (Niwa et al., 2005; Ralston et al., 2010; Tanaka et al., 1998). However, the properties of the two cell line systems are not exactly the same (Cambuli et al., 2014) and both are likely to be different from the embryonic trophectoderm. To test this hypothesis, we analyzed global gene expression pattern from the different cell sources.

We utilized the inducible *Cdx2* over-expression ES cell system (Niwa et al., 2005; Ralston et al., 2010) to measure transcriptome changes upon single gene perturbation. Time-course microarray analysis was performed on three different inducible clones at day 0, day 0.25, day 1, day 2 as well as day 6. Changes in individual gene expression during the time-course are shown in Figure 1A. CDX2-induced gene activation or repression may start as early as 6 hours after over-expression. On day 6, the TE transcriptional program (including *Cdx2, Tcfap2c* and *Id2*) is fully activated, while the ES transcriptional program (including *Pou5f1, Sox2* and *Nanog*) is completely repressed. Notably, a list of genes including *Hoxa9, Hoxa10, Hoxb6, Foxh1, Phf19, Nkx1-2* and *Sox7* is transiently induced during the early time points, but eventually repressed on day 6. As the chromatin state of ES cells is relatively open, forced expression of *Cdx2* may activate targets that are irrelevant to trophectoderm development.

**Figure 1.**
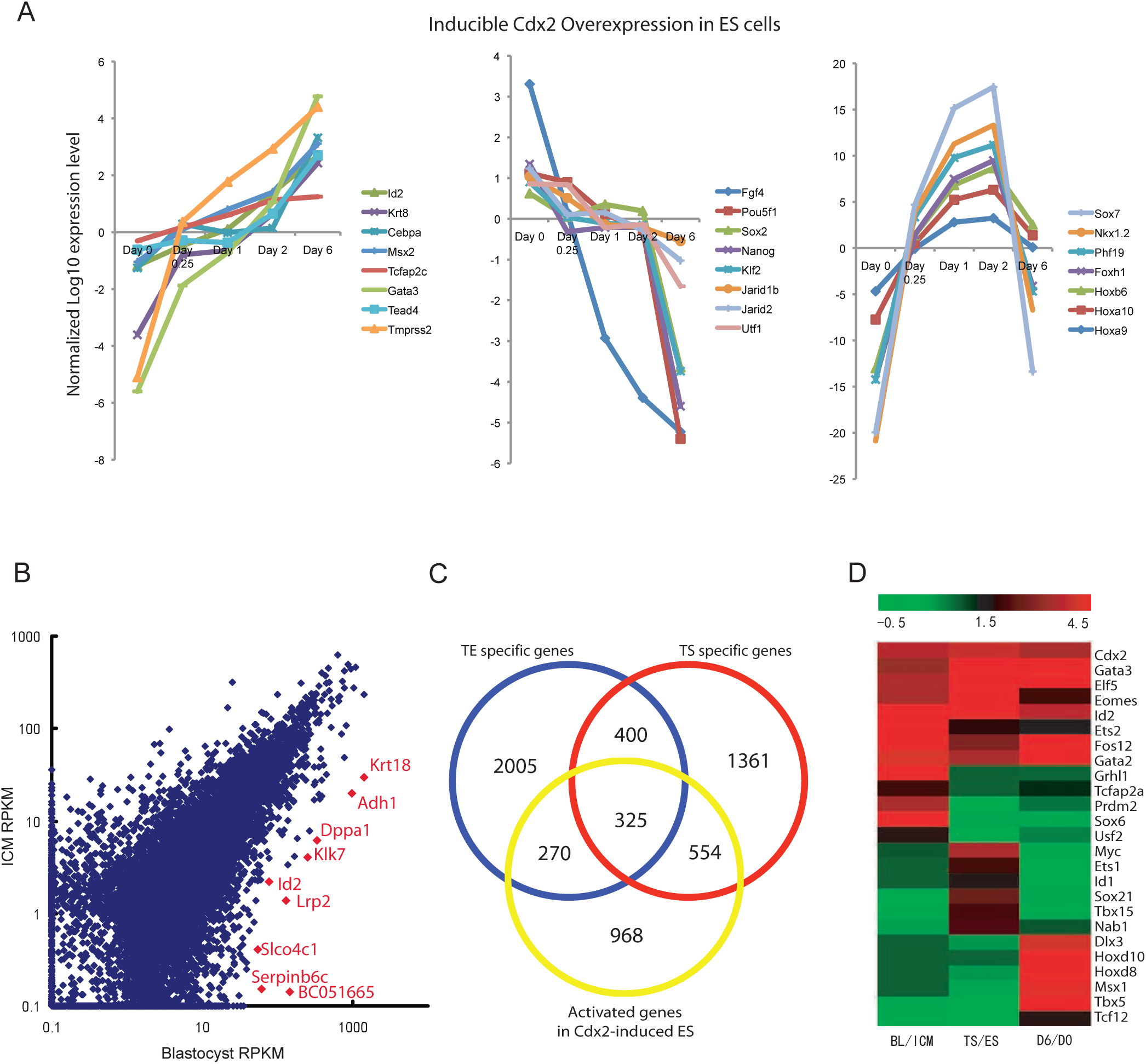
Comparison of expression profiles from different trophoblast cellular systems. (A)*Cdx2* over-expression in ES cells induces trophoblast differentiation. The plot depicts gene expression changes of selected genes (average in three inducible *Cdx2* over-expressing ES clones) during the differentiation time course. (B)A scatter plot to compare gene RPKM values in the blastocyst RNA samples and the ICM RNA samples. Examples of TE specific markers identified by mRNA-seq analysis are highlighted in red. (C)Comparison of TE specific gene list (from mRNA-seq analysis of the blastocyst tissues), TS specific gene list (from microarray profiles of TS cells compared to ES cells, Kidder and Palmer, 2010) and Cdx2 OE upregulated gene list (from microarray profiles of Day 6 Cdx2 over-expression compared to Day 0 un-induced ES cells). (D)Gene expression heat map comparing lineage specific markers in different trophoblast systems. *Grhl1* and *Tcfap2a* are high in the trophectoderm tissues, while TS cell shows higher *Myc* expression. In addition, Cdx2 induces *Hox* gene expression in ES cells. BL, blastocyst.

In order to generate the whole-genome gene expression profiles of *in vivo* TE, we extracted RNA from E3.75 blastocysts as well as E3.75 ICMs isolated by immuno-surgery. Global mRNAs were then amplified against poly-A tail and analyzed by high throughput sequencing following the single cell mRNA-seq protocol (Tang et al., 2009). Using a modified version of BioScope software, we have calculated the RPKM value (reads per kilobase of exon model per million mapped reads) for each gene as a measurement of expression level quantification. In the E3.75 blastocyst, the majority of the cells are TE cells; TE specific genes should be only detected in the whole blastocyst samples but not in the ICM samples. A comparison of individual gene mRNA-seq RPKM value between the two samples reveals the TE/ICM differential expressions (Figure 1B, and Table S1). We sorted genes by their expression fold difference between whole blastocysts and ICMs; and then define the top 3000 genes as TE specific markers. *Id2* and *Dppa1*, which have already been characterized in our previous single cell based study (Guo et al., 2010), are among the top of the TE specific genes as shown in the scatter plot (Figure 1B). This data set provides comprehensive information about *in vivo* gene expression patterns in the two segregated blastocyst cell lineages.

In addition, we compared *in vitro* TS and ES gene expression profiles and generated TS specific gene list from the published microarray data (Kidder and Palmer, 2010; Rugg-Gunn et al., 2010). We then identified genes that are significantly higher in the Day 6 *Cdx2* over-expressed ES cells compared to un-induced ES cell control. When comparing these data, we found lineage-specific expression patterns differ between *in vitro* culture systems and the *in vivo* embryonic tissues (Table S1). In addition, the TE signature has a higher overlap with TS cells compared to *Cdx2* over-expressing ES cells, consistent with Hemberger’s previous study (Cambuli et al., 2014; Latos et al., 2015) (Figure 1C). As shown in Figure 1D, although the three systems (BL/ICM, TS/ES, D6/D0) share common lineage specific markers such as *Cdx2, Eomes, Gata3, Elf5* and *Id2*, they possess distinct transcriptional programs: *Tcfap2a* and *Grhl1* expressions are high in the trophectoderm, while Myc and Id1 expressions are high in the TS cells. In particular, our time course analysis with the *Cdx2* over-expressing ES cells suggests that CDX2 activates the Hox gene clusters. ChIP-seq data by Nishiyama et al. (2009) confirmed CDX2 binding to different Hox genes in the ES cell system. However, the vast majority of these Hox genes are not expressed in the TE tissues according to our RNA-seq data. Although *Hox* genes are potential CDX2 targets in the developing embryo itself (Lohnes, 2003), their detection here is likely not functionally meaningful during the trophoblast lineage development, consistent with the observation that their expression is not maintained in TS cells.

### Identification of CDX2 functional targets in *Cdx2* knockout embryos

We next used *Cdx2* knockout embryos to identify genes whose expression level depends on CDX2. Cdx2 is first activated at E2.5 at 8-cell stage. Cdx2 null embryos die at around E4.5: they still form a blastocoel but fail to maintain blastocyst integrity (Ralston and Rossant, 2008; Strumpf et al., 2005). In order to characterize *Cdx2* functional targets *in vivo*, we applied high throughput microfluidic qPCR gene expression profiling on E3.75 blastocysts from *Cdx2+/−* crosses. In total, 27 blastocysts were assayed against 176 genes selected partly by our RNA-seq data (Data not shown). 6 out of 27 embryos do not have any detectable *Cdx2* mRNA (Figure 2A, and Table S2). These embryos also have significantly higher Neo expression, which was used to replace the *Cdx2* alleles. These embryos were designated as Cdx2-/- embryos. In addition, 4 embryos had negligible Neo levels: these were designated as wild-type embryos. As shown in Figure 2A, expression of *Pou5f1, Nanog* and *Sox2* in the 6 *Cdx2* null embryos were upregulated comparing to the heterozygous and wild-type, suggesting that CDX2 is required to repress ES pluripotent markers during early embryogenesis. The expression of the TE marker *Gata3* is unperturbed, consistent with a previous study demonstrating that Cdx2 regulates trophoblast development in parallel to Gata3 (Ralston et al., 2010). In total, more than 50 percent of all assayed genes show reproducibly altered gene expression levels in the *Cdx2* null blastocysts. The hierarchical clustering of gene expression profiles of the 27 blastocysts clearly demonstrated that the 6 *Cdx2* knockout blastocysts exhibited distinct global gene expression patterns (Figure 2B).

**Figure 2.**
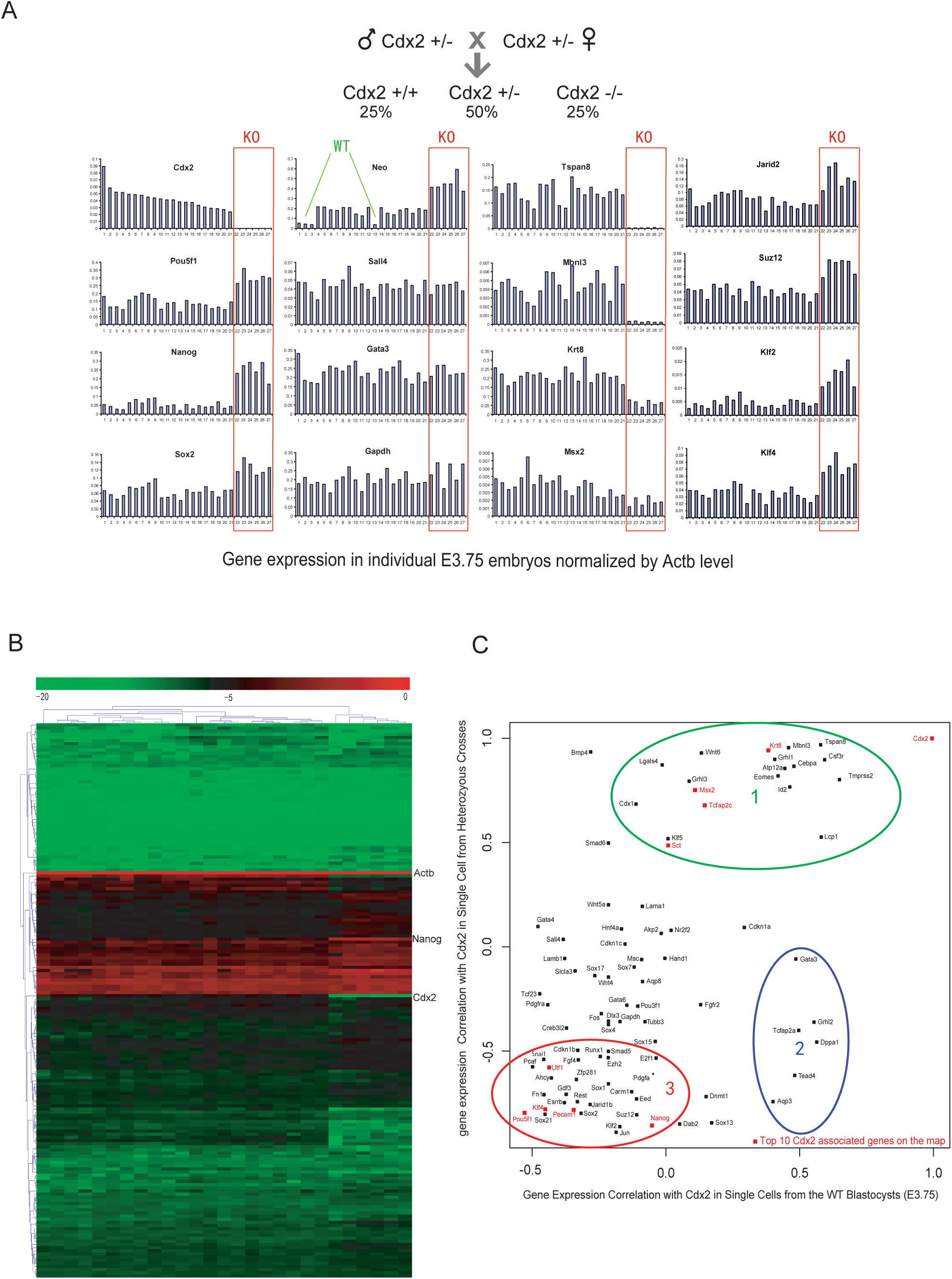
Identification of Cdx2 functional targets in vivo from E3.5 Knockout blastocysts.

(A)qPCR analysis of E3.75 blastocysts derived from *Cdx2* heterozygous intercrosses. Each bars represents one blastocyst. All expression levels are normalized against endogenous control *Actb*. The order of the embryos is sorted according to *Cdx2* expression.

(B)Hierarchical clustering of expression profiles of all analyzed individual blastocysts.

(C)Expression correlation map of different genes to *Cdx2*. X-axis indicates gene correlation with *Cdx2* in single cells harvested from ∼E3.75 wild type embryos. Y-axis indicates gene correlation with *Cdx2* in E3.75 blastocysts harvested from *Cdx2* +/− intercrosses. See text for discussion of cluster 1, 2, and 3.

In order to reveal the hierarchy of CDX2-regulated gene expression, we generated single cell expression profiling data from the wild-type E3.75 blastocyst using our previous methods (Guo et al., 2010) and plotted genes according to their correlation with *Cdx2* in both single cell data and the knockout blastocyst data (Figure 3C). Based on the 2D plot, there are mainly three groups of genes that are of interest. The first group of genes are highly correlated with *Cdx2* expression and thus can be confirmed as being positively regulated by CDX2. The second group of genes are independent of *Cdx2* activation, however they are specific to the CDX2-positive TE, which indicates they function in parallel with *Cdx2*. Finally, the third cluster genes include most of ICM pluripotent markers that are negatively regulated by *Cdx2*. Our results clearly demonstrate that *Cdx2* activates the TE transcriptional program and represses the pluripotent network during blastocyst formation.

**Figure 3.**
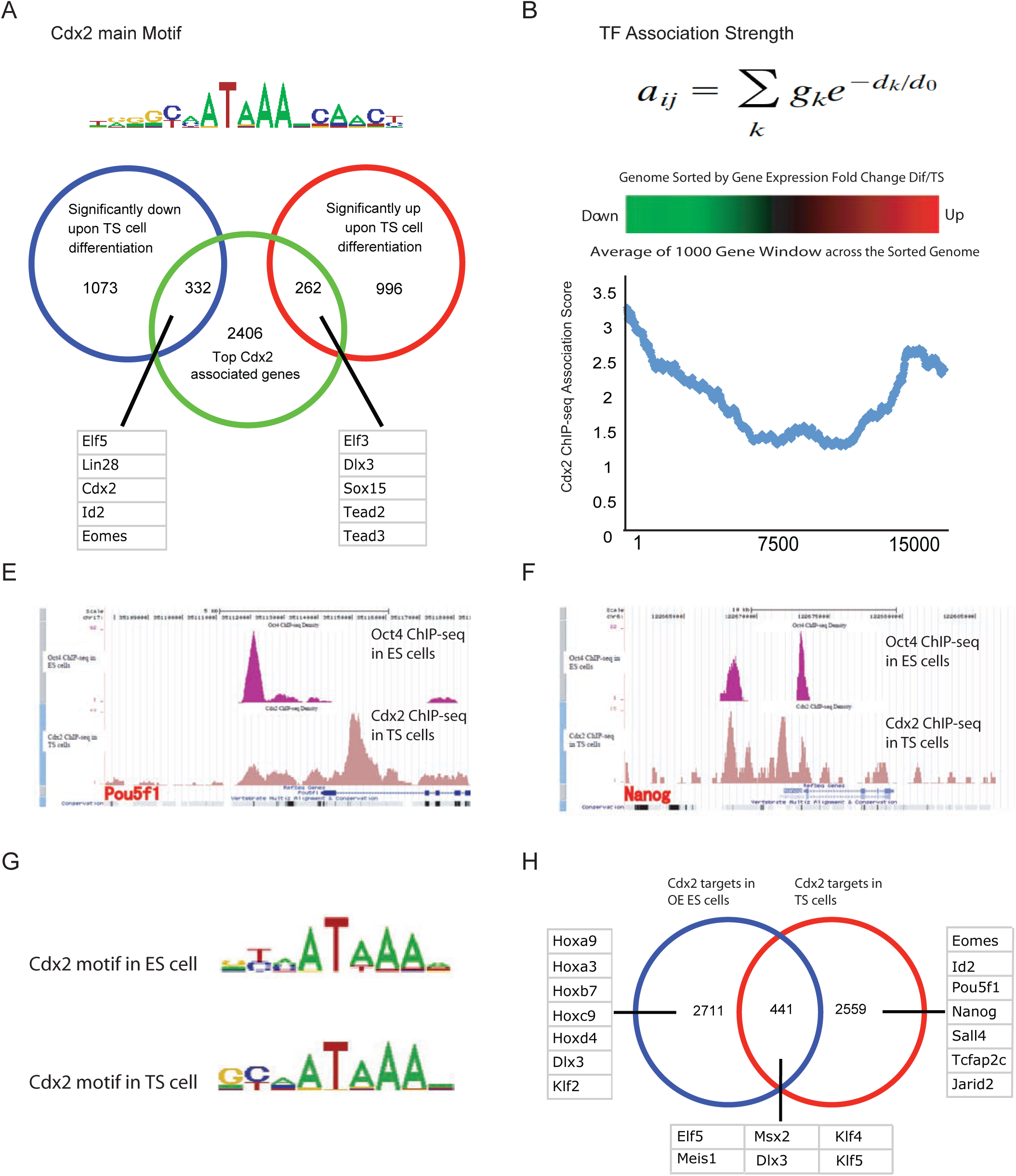
Cdx2 ChIP-Seq analysis in TS cells reveals direct targets of Cdx2 (See also Figure S1) (A)Cdx2 main binding motif clusters identified with CisFinder via 200bp sequences centered at ChIP sites. (B)The scoring system of transcription factor ChIP-Seq binding association. (C)Venn diagram showing the overlap between Cdx2 target list and the significantly up/down regulated genes after 6 days of TS cell differentiation. Representative Cdx2 targets are listed. (D)Blue line: relationship between gene expression difference and TF ChIP-Seq association score. X-axis shows the gene rank after sorting the genome according to expression fold change between differentiated and undifferentiated TS cells (Kidder and Palmer, 2010). Y-axis shows the average Cdx2 binding association score from a sliding window of 1000 genes. (E)Oct4 ChIP-Seq peaks (from ES cells) and Cdx2 ChIP-Seq peaks (from TS cells) in the Pou5f1 gene region viewed with USCS mouse mm8 browser. (F)Oct4 ChIP-Seq peaks (from ES cells) and Cdx2 ChIP-Seq peaks (from TS cells) in the Nanog gene region viewed with USCS mouse mm8 browser. (G)Analysis of Cdx2 ChIP-seq results from our TS cell system and the ES cell Cdx2 overexpression system (Nishiyama et al., 2009) reveals strikingly similar core Cdx2 binding motifs. (H)Although Cdx2 does not bind to Pou5f1, Sox2 and Nanog in the ES cell TE differentiation system (Nishiyama et al., 2009), we have observed significant repressive bindings of Cdx2 on pluripotent genes in the established TS cell system.

### Whole-genome ChIP-Seq analysis reveals diverse targets of CDX2 in TS cells

To characterize genome-wide direct targets of CDX2, we performed chromatin immunoprecipitation sequenceing (ChIP-Seq) experiment within TS cells using a highly specific CDX2 antibody (Figure S1A). Enriched CDX2 binding DNA fragments were analyzed by high-throughput sequencing. Using model-based ChIP-seq analysis (Zhang et al., 2008), we found a total of 16736 confident peaks (Table S3). CDX2 mainly binds within the window from 1kb upstream to 1kb downstream of target transcriptional start sites (Figure S1B). We performed de novo motif discovery with CisFinder (Sharov and Ko, 2009). The main consensus-binding motif cluster turns out to be a known CDX motif (Figure 3A). This motif is overrepresented in ChIP-enriched regions, as we looked at motif counts across 4kb windows centered on the CDX2 binding sites (Figure S1C). Traditional ChIP-Seq analysis usually associates a TF binding peak with a gene based on the distance between the peak and the transcriptional start site of the gene. Ouyang et al. (Ouyang et al., 2009) made a significant improvement by integrating the surrounding peak intensity and the proximity to genes to define the association strength between TF and individual genes (Figure 3B). We used this method to calculate CDX2 ChIP-seq association score for each gene, ranked the genome according to the association score and defined top 3000 associated genes as CDX2 targets (Table S1). To associate CDX2 bindings with its function in TS cells, we reanalyzed published gene expression data for TS cells as well as differentiated TS cells (Kidder and Palmer, 2010). We identified genes that are significantly up- or down-regulated during 6 days of TS cell differentiation (Table S1). *Cdx2* was dramatically down-regulated upon TS differentiation, together with other TS cell markers such as *Eomes* and *Elf5*, as previously shown (Kidder and Palmer, 2010). We overlapped our defined CDX2 targets with the significantly up or down-regulated genes upon TS cell differentiation. Among the TS cells specific genes, CDX2 binding associated genes include *Elf5, Lin28, Cdx2, Id2* and *Eomes* (Figure 3C). Potential CDX2 negative targets such as *Elf3, Dlx3* and *Sox15* (Differentiation markers) also have high CDX2 binding association. We sorted the genome according to expression fold difference between differentiated and undifferentiated TS cells. We then looked for changes in the average association score of CDX2 in a sliding 1000 gene window. Interestingly, TS stem cell signatures or potential CDX2 positive targets have the highest association score (Figure 3D). Genes whose expression did not change during differentiation have extremely low association scores. However, TS differentiation markers or potential CDX2 negative targets tend to have moderate CDX2 binding. The CDX2 binding association curve suggests that CDX2 is actively involved in both gene activation and repression within the TS cells.

Previous studies have revealed that CDX2 is also involved in ICM/TE lineage segregation by repressing ES core pluripotent gene Oct4 (*Pou5f1*) (Niwa et al., 2005; Strumpf et al., 2005). Early studies have employed the *Cdx2*-overexpressing ES cell system to demonstrate that CDX2 directly interacts with OCT4 to repress its transcriptional activity in ES cells. ChIP-qPCR assay showed that CDX2 prevented OCT4 protein from binding to an auto-regulatory element (ARE) of *Pou5f1* (Niwa et al., 2005), but CDX2 did not whereas CDX2 directly bind to *Pou5f1* and *Nanog* regulatory elements during the initial phases of CDX2 overexpression in ES cells (Nishiyama et al., 2009a). In our CDX2-ChIP-Seq experiments on TS cells, however, we found clear and significant CDX2 binding to cis-acting sequences associated with the core ES transcription factors: *Pou5f1* and *Nanog* (Figure 3E and 3F). CDX2 binds to the first intron of *Pou5f1* with more than 40-fold enrichment, validated by ChIP-qPCR across the binding region (Figure S1D). In addition, it is noteworthy that CDX2 binds to Pou5f1 at the first intron instead of promoter or other cis-regulatory DNA elements of *Pou5f1*. The exact role of this intronic region in regulating *Pou5f1* expression remains to be explored.

To identify more genes whose expression is regulated by CDX2 binding, we compared CDX2 ChIP-seq data from TS cells and the CDX2 overexpression ES system from published data (Nishiyama et al., 2009a). Interestingly, although the core CDX2 binding motifs are remarkably the same (Figure 3G), the actual targets vary in the two systems (Figure 3H). Contrary to our CDX2-ChIP-Seq data in TS cells, CDX2 does not bind to the core pluripotent genes, *Sox2, Pou5f1* and *Nanog*, during the initial step of ES cell differentiation upon CDX2 over-expression. However, binding was observed in association with several *Klf* genes. As our data show in Figure 1D, the gene expression patterns and regulatory network in the CDX2 overexpressing ES cell system are not exactly identical with those in the TS cell system. Because of this, we used ES and TS cell comparisons to define CDX2 regulatory activity in trophoblast development in the rest of the studies.

### Cdx2 as a lineage-specific transcriptional repressor during ICM/TE segregation

To compare CDX2 and OCT4 binding sites, we integrated OCT4-ChIP-Seq data in ES cells (Whyte et al., 2013) with our TS data. By evaluating the genome association score (Figure 3B), we defined the top 3000 OCT4 binding targets in ES cells (Table S1). Remarkably, OCT4 and CDX2 have a significant overlap in downstream targets (Figure S2A). Binding site overlap studies also revealed 449 co-occupied loci. Many important pluripotent markers such as *Nanog, Sox2, Klf4, Esrrb* and *Utf1* are within the overlapping targets list (Figure S2B). To associate CDX2 and OCT4 binding with function in TS cells and ES cells respectively, we overlapped the top 3000 CDX2 binding targets with TS-specific genes and ES-specific genes, and did the same with OCT4 binding targets. Venn diagrams reveal that both CDX2 and OCT4 not only bind to TS-expressed genes, but also to ES-expressed genes (Figure S2C and S2D). Notably, ES cell-specific genes *Jarid2* and *Nanog* are among the top 5 genes bound by both CDX2 and OCT4, suggesting that CDX2 and OCT4 play active, but opposite, roles in regulating *Jarid2* and *Nanog* expression. Furthermore, we sorted the genome according to the expression fold difference between TS and ES cells and looked for change in the average association score of CDX2 in a sliding 1000 gene window. Interestingly, gene activation or potential positive targets have the strongest binding association (Figure 4A), whereas gene repression or potential negative targets tend to have moderate binding association. Similar patterns were also found for OCT4 (Figure 4B). The significance of expression fold change between TS and ES cells is positively correlated with binding association score regardless of whether it is gene activation or repression.

**Figure 4.**
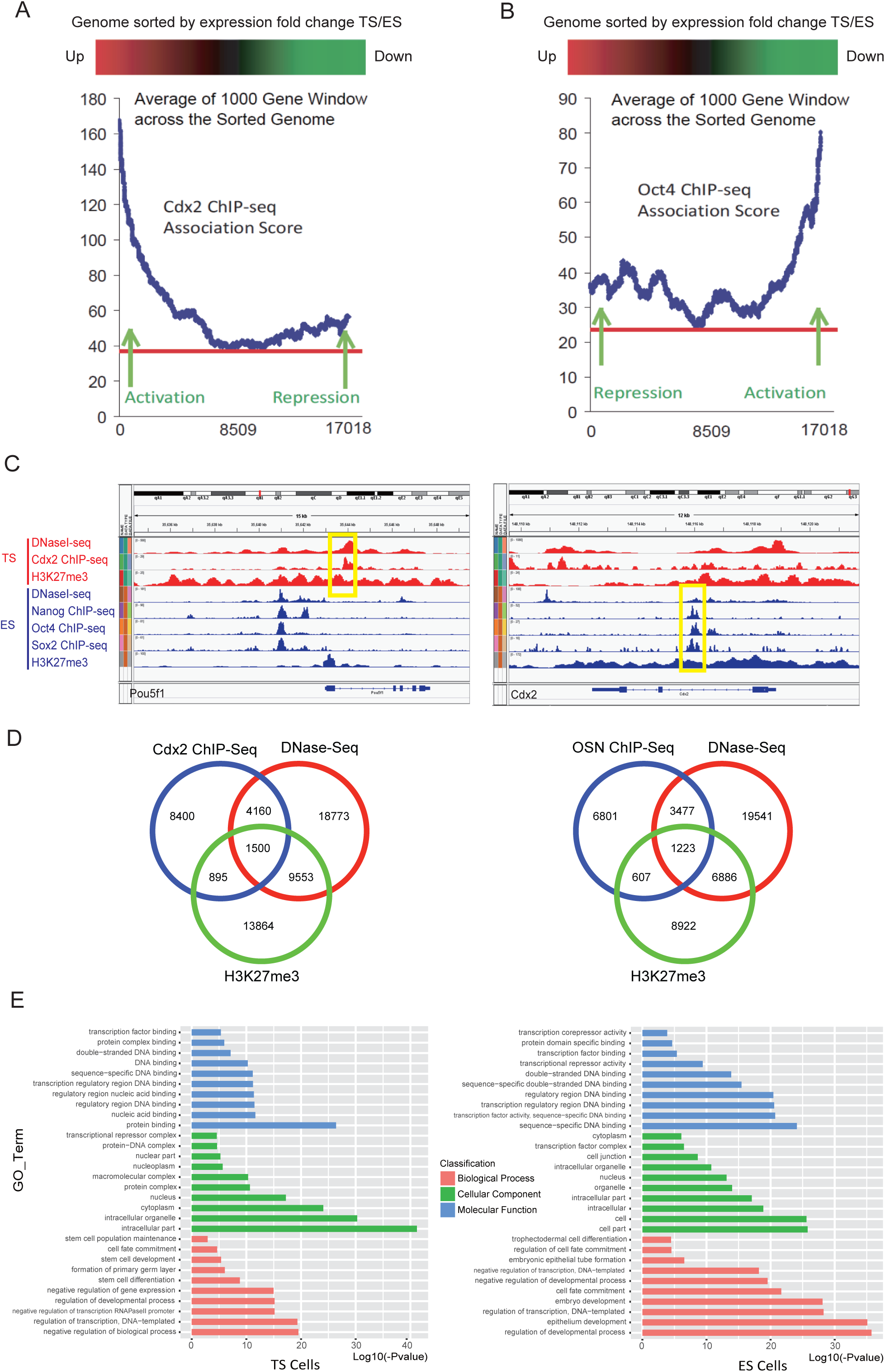
Cdx2 directly competes with Oct4 on genome-wide regulation of lineage segregation (See also Figure S2 and S3) (A)Relationship between gene expression difference of TS/ES and Cdx2-ChIP-Seq association score. X-axis shows the gene rank after sorting the genome according to expression fold change between TS and ES cells. Y-axis shows the average Cdx2 binding association score from a sliding window of 1000 genes. (B)Relationship between gene expression difference of TS/ES and Oct4-ChIP-Seq association score. X-axis shows the gene rank after sorting the genome according to expression fold change between TS and ES cells (Kidder and Palmer, 2010). Y-axis shows the average Oct4 binding association score from a sliding window of 1000 genes. (C)Cdx2 ChIP-Seq peaks, H3K27me3 Peaks, DNase Peaks from TS cells in the *Pou5f1* gene region viewed with IGV; OSN ChIP-Seq peaks, H3K27me3 Peaks, DNase Peaks from ES cells in the Cdx2 gene region viewed with IGV. OSN: Oct4-Sox2-Nanog. (D)Venn diagram show silencer candidates in TS cell (left); Venn diagram show silencer candidates in ES cell (right). (E)GO analysis of silencer-related genes in TS cells; GO analysis of silencer-related genes in ES cells.

Our ChIP-Seq data indicates that CDX2 extensively binds and represses other ES-specific genes in addition to *Pou5f1* in TS cells. Therefore, how CDX2 binding leads to transcriptional repression instead of activation becomes a key question. Mapping DNase I hypersensitive sites (DHSs) by DNase-Seq has been a valuable tool for identifying different types of regulatory DNA elements, including promoters, enhancers, and silencers (Boyle et al., 2008). Here, we integrated published DNase-Seq data in TS and ES cells, respectively (Calabrese et al., 2012). Results show that DHSs and CDX2 repressive binding sites co-occupy the first intron of *Pou5f1* in TS cells (Figure 4C). Similarly, the CDX2 binding sites at other ES pluripotent markers such as *Nanog, Sox2* and *Klf4*, are also DNase I hypersensitive sites (Figure S3A). Previous study has shown that core transcription factors OCT4, SOX2, and NANOG share substantial target genes (Young, 2011) and therefore we further analyzed OCT4, SOX2, and NANOG ChIP-Seq, plus DNase-Seq in ES cells (Whyte et al., 2013). Interestingly, the three transcription factors co-occupy a *Cdx2* intron with high DNase I hypersensitive signal (Figure 4C). Similar binding patterns also exist in other TS-specific genes, including *Id2, Tcfap2a*, and *Msx2* (Figure S3B). Notably, there are high level of H3K27me3 signals surrounding these binding sites, indicating that the regions are transcriptionally inactive (Li et al., 2007). Together, this type of binding site can be considered as lineage-specific silencers, based on their unique chromatin state and lineage-specific transcriptional repression features.

To characterize lineage-specific silencers on a genome-wide scale, we defined silencer candidates in TS cells based on possession of three properties: CDX2 binding, DNase I hypersensitivity, and enriched H3K27me3. This identified 1500 putative CDX2-regulated TS-specific silencers in TS cells (Figure 4D, and Table S4). Similarly, we defined silencer candidates in ES cells as sharing Oct4-Sox2-Nanog (OSN) co-binding peak, DNase I hypersensitive sites, and enriched H3K27me3. This produced a list of 1223 putative OSN-regulated silencer regions in ES cells (Figure 4D, and Table S4). Candidate silencer elements were annotated to the nearest TSS and its associated gene. GO analysis revealed that top associated biological process of silencer-related genes in TS cells includes “stem cell differentiation”, “formation of primary germ layer” and “cell fate commitment”, which suggests that majority of CDX2-repressed genes in TS cell are pluripotent genes that function in early embryogenesis (Figure 4E, and Table S4). Conversely, the top associated biological process of silencer-related genes in ES cells includes “trophectodermal cell differentiation”, indicating that OSN may repress TS-specific genes in ES cells through binding their surrounding silencers (Figure 4E, and Table S4).

## DISCUSSION

In this study, we use single cell qPCR and RNAseq on normal and *Cdx2* knockout embryos to provide new insights into the likely downstream targets of *Cdx2* function in the developing blastocyst.In *Cdx2* knockout embryos, a large portion of ICM markers are up-regulated while TE markers are down-regulated, demonstrating that *Cdx2* is a key regulator, playing a dual function in TE formation through both repressing pluripotency genes and activating TE genes.

To investigate the direct targets of CDX2 in the whole genome, we performed CDX2 ChIP-Seq in TS cells. It turns out that CDX2 binds to a wide range of targets, including both markers for TS self-renewal and differentiated state. Furthermore, we integrated CDX2 function with other TS specific transcription factors and compared our CDX2 target list with the published EOMES and TCFAP2C target lists in TS cell (Kidder and Palmer, 2010)(Table S1). We showed that CDX2, EOMES and TCFAP2C co-occupy 314 genes (Figure S4A). On the top of this overlapping target list, there are many characterized TE lineage markers such as *Id2, Elf5* and *Hand1*. Interestingly, the top overlapping targets of CDX2, EOMES and TCFAP2C are TE-specific activation targets. In the ChIP-seq data from the established TS cultures, we also observed CDX2 binding to many important pluripotent genes such as *Pou5f1, Nanog, Sox2* and *Esrrb*, genes that are also upregulated in *Cdx2* mutant blastocysts. This might suggest that direct repressive occupancy by CDX2 is involved in maintaining repression of pluripotent gene expression in the trophoblast lineage. By contrast, in the *Cdx2*-inducible overexpression ES system, it had previously been shown that CDX2 does not directly bind to ES cell core regulatory genes ((Nishiyama et al., 2009b). However, this system may be more representative of the early transient phases of lineage activation, rather than the stable regulatory system needed for TS cell lineage maintenance.

A subset of CDX2 binding sites in TS cells possess a unique chromatin state, with DNase I hypersensitivity, indicative of open chromatin, and enrichment of the H3K27me3 modification. These sites are enriched in associations with known pluripotency regulatory genes and are proposed to function as lineage specific silencers mediating transcriptional repression of pluripotent markers (*Pou5f1, Sox2* and *Nanog*) to maintain TS cell identity. This is consistent with *in vivo* studies showing that *Cdx2* knockout embryos fail to down-regulate *Pou5f1* and *Nanog* in the trophectoderm (Strumpf et al., 2005). Strikingly, re-analyzing the published OSN ChIP-Seq data in ES cells reveals that the ES cell core transcription factors, OSN, co-occupy regulatory regions of TS-specific genes with DNase I hypersensitive feature and enriched H3K27me3 modification, similar to CDX2 targets in TS cells. Most work on ES cell gene regulatory networks has focused on defining enhancers that function in activating the ES cell pluripotent program. However, potential silencing in ES cells defined by our study offers a new perspective to broaden the understanding of the pluripotent state as one in which both activation of epiblast gene expression and inhibition of extra-embryonic gene expression is needed for stability of the stem cell state. Future studies such as identification of co-activators or partners binding to silencers are needed to reveal the molecular mechanisms underlying how candidate silencers function in the process of transcriptional repression. Importantly, our study provides valuable resources to study the function of silencers, one of the cis-regulatory DNA elements, in regulating gene transcription in diverse biological processes.

In conclusion, our findings provide evidence that lineage-specific silencers exist in both TS and ES cells on a genome-wide scale. We identify *Cdx2* as lineage-specific repressor that transcriptionally represses ICM pluripotency genes via binding to specific silencer elements. Finally, we propose that lineage-specific transcriptional repression through silencers may serve as a novel mechanism to establish TS/ES unique gene expression patterns, and promote ICM and TE lineage segregation during first cell fate determination.

## EXPERIMENTAL PROCEDURES

### Culture of ES cells and TS cells

ES cells were maintained in Dulbecco’s modified Eagle’s medium (DMEM, Gibco-BRL), with 20% heat-inactivated ES-qualified fetal bovine serum (FBS, Gibco-BRL), 0.055 mM □-mercaptoethanol (Gibco-BRL), 2mM L-glutamine, 0.1 mM MEM nonessential amino acid, 5000 U/ml penicillin/streptomycin and 1000 U/ml leukemia inhibitory factor (LIF, Chemicon) without MEFs. TS cells are maintained in DMEM, 20% of FBS, 0.1 mM sodium pyruvate, 0.1 mM nonessential amino acids, 0.055 mM □-mercaptoethanol, 2 mg/ml of sodium heparin (Sigma), and 20 ng/ml of recombinant FGF4 (Sigma) in the presence of 70% of the MEF-conditioned medium.

### RNA extraction and gene expression microarray

Cells were rinsed twice in ice-cold PBS. Total RNA was extracted using Trizol (Invitrogen) and column-purified with the RNeasy kit (Qiagen). The quality of RNA was then checked by agilent 2100 bioanalyzer. High quality samples are analyzed using the MouseRef-8 v2 Expression BeadChip (Illumina).

### mRNA-seq analysis of individual blastocysts and dissected ICMs

E3.75 ICMs were collected by immnosurgery (Solter and Knowles, 1975). Total RNA from blastocysts or ICMs was extracted using the PicoPure RNA isolation kit (Arcturus Bioscience), and then reverse transcribed using polyT primers. Terminal deoxynucleotidyl transferase was used to add a polyA tail to the 3 end of first-strand cDNAs, which was followed by PCR amplification of the cDNAs. After shearing into 80-130 bp fragments, P1 and P2 adaptors were ligated to each end, and the fragments were subjected to 8-10 cycles of PCR amplification. Amplified samples are sequenced using SOLiD sequencer (Applied Biosystems). ABI’s whole transcriptome software tools were used to analyze the sequencing reads. The reads obtained were matched to the Mouse genome (mm 9). Feature counts were normalized using the RPM (read per million aligned reads) method.

### Gene expression profiling of individual blastocysts by high throughput microfluidic qPCR

Total RNA was extracted from individual mouse embryos using the PicoPure RNA isolation kit (Arcturus Bioscience). The entire RNA preparation was used for cDNA synthesis at 37°C for 2 hrs using the high capacity cDNA archive kit (Applied Biosystems). One eighth of each cDNA preparation was pre-amplified using the TaqMan primers for genes of interest by 16 cycles of amplification (each cycle: 95°C for 15 Sec and 60°C for 4 min) using the TaqMan PreAmp Master Mix Kit (Applied Biosystems). These preamplified products were diluted 5-fold before analysis. Real-time reactions were performed in technical triplicate with master mix (Applied Biosystems) in 48.48 Dynamic Arrays on a BioMark System (Fluidigm). Threshold cycle (Ct) values were calculated from the system’s software (BioMark Real-time PCR Analysis) and used as a direct measure of gene expression.

### High throughput single cell qPCR

Equal volumes of each inventoried TaqMan Gene Expression Assay (20X, Applied Biosystem) were pooled and then diluted using TE buffer so that each assay was at a final concentration of 0.2X. These pooled assays were for use in the pre-amplification step. Individual cells were harvested directly into the 10 μL RT-PreAmp Master Mix (5.0 μL CellsDirect 2X Reaction Mix (CellsDirect qRT-PCR kit, Invitrogen); 2.5 μL 0.2X Assay Pool; 0.2 μL RT/Taq Enzyme (CellsDirect qRT-PCR kit, Invitrogen); 2.3 μL Rnase-free water. The harvested single cell samples were immediately frozen and stored at -80°C. Cell lysis and sequence-specific reverse transcription were performed at 50°C for 20 min. The reverse transcriptase was inactivated by heating to 95°C for 2 min. Subsequently, in the same tube, cDNA went through sequence-specific amplification by denaturing at 95°C for 15 s, and annealing at 60°C for 4 min for 18 cycles. The pre-amplified products were diluted 5-fold and then analyzed by TaqMan PCR. Real-time reactions were performed with Universal PCR Master Mix and inventoried TaqMan gene expression assays (Applied Biosystems) in 48.48 Dynamic Arrays on a BioMark System (Fluidigm). Threshold cycle (Ct) values were calculated from the system’s software (BioMark Real-time PCR Analysis, Fluidigm).

### Cdx2 ChIP-Seq with TS cells

TS cells were cross-linked with 1% (w/v) formaldehyde for 10 min at room temperature, and formaldehyde was then inactivated by the addition of 125 mM glycine. Chromatin extracts containing DNA fragments with an average size of 500bp were immunoprecipitated, using anti-Cdx2 (CDS-88, Biogenex). The ChIP enriched DNA was then decross-linked and sequenced by Genome Analyzer II (Illumina) according to Illumina’s manuals. Peak calling based on the Cdx2 ChIP-seq data was performed using MACS.

### ChIP-seq data analysis

De novo motif discovery was performed with CisFinder. The association strength between TF and individual genes were calculated with the published method (Ouyang et al., 2009). The association score pattern along the gene expression change window was generated using EXCEL. Briefly, genes were ordered according to their expression fold change between the compared samples, then the average association scores in the sliding 1000 gene window were cacluated across the sorted genome. ChIP-seq and DNaseI-seq peaks were visualized with IGV software. Overlapping motif analysis was performed using CDX2 ChIP-seq data and previously published OCT4 ChIP-seq data (Whyte et al., 2013). We allowed up to 200 bp between the borders of two peaks.

## AUTHOR CONTRIBUTIONS

Experiments were designed by G.G., carried out by G.G., X.H., P.Y., A.R., L.S., M.H., T.M., L.P., G.Y. and J.J., analyzed by D.H., G.G. The paper was written by D.H, J.R., P.R. and G.G. This work was supported by Zhejiang University Stem Cell Institute.

## ACKNOWLEDGMENTS

We thank Y. Zhou, L. Shen, S. Ying and D. Zhou for help on experiments. We thank Y. Hong, H. NG and C. Neil for insightful suggestions for the project. This work was supported by funding from Fundamental Research Funds for the Central Universities (G.G.) and 1000 Youth Talent Plan (G.G.). G.G. is a professor from Zhejiang University.

